# Multipad Agarose Plate (MAP): A Rapid and High-Throughput Approach for Antibiotic Susceptibility Testing

**DOI:** 10.1101/2024.01.20.576355

**Authors:** Morten Kals, Leonardo Mancini, Jurij Kotar, Allen Donald, Pietro Cicuta

## Abstract

We describe a phenotypic antibiotic susceptibility testing (AST) method that can provide an eightfold speedup in turnaround time compared to the current clinical standard by leveraging advances in microscopy and single-cell imaging. A newly developed growth plate containing 96 agarose pads, termed the Multipad Agarose Plate (MAP), can be assembled at low costs. Pads can be prepared with dilution series of antibiotics. Bacteria are seeded on the pads and automatically imaged using brightfield microscopy, with a fully automated segmentation pipeline quantifying microcolony formation and growth rate. Using a test set of nine antibiotics with very different targets, we demonstrate that accurate minimum inhibitory concentration (MIC) measurements can be performed based on the growth rate of microcolonies within three hours of incubation with the antibiotic. Faster, reliable and high throughput methods for AST, such as MAP, could improve patient care by expediting treatment initiation and alleviating the burden of antimicrobial resistance.

## 1 Introduction

Antibiotic resistance is a growing issue in healthcare worldwide. In 2019, antibiotic resistance was classified as the primary cause of death for over 1.3 million people worldwide, and this number is projected to rise to over 10 million annual deaths by 2050 [1]. Resistance is accelerated by antibiotic overuse and misuse [2]. To limit the incorrect prescription of antibiotics, it is necessary to have fast and accurate methods for antibiotic susceptibility testing (AST), where the effectiveness of antibiotics against a particular bacterial strain is determined [1]. AST can be performed using either genotypic or phenotypic methods.

Genotypic AST identifies specific genetic elements associated with resistance to certain antibiotics, such as resistance genes or mutations. These methods are fast and can predict resistance even before it becomes phenotypically apparent. However, they may not account for all resistance mechanisms as not all resistance genes are known or detectable, leading to systematic false negatives [3]. In addition, non-genetic resistance mechanisms, such as epigenetic changes or phenotypic heterogeneity, are not directly detectable. Epigenetic changes, which alter gene expression patterns without causing DNA mutations, can inheritably alter gene expression and are known to contribute to antibiotic resistance [4]. Phenotypic heterogeneity, which originates from stochastic changes in microbial physiology [5], is also known to play a major role in the development of resistance, for example via the production of persister cells [6]. These mechanisms can allow bacteria to survive antibiotic treatment without any changes in their genetic makeup [7].

Phenotypic AST, on the other hand, directly measures the growth of bacteria in the presence of antibiotics, thereby detecting whether the bacteria are susceptible or resistant to specific drugs. Current phenotypic methods include disk diffusion, ETest strips and broth microdilution [8]. This family of methods can detect all types of resistance mechanisms, known or unknown, as the organism’s actual response to an antibiotic is measured. However, these methods are typically slower than genotypic testing, typically requiring 16-48 hours for bacterial growth [9, 10].

Clinical studies have shown that hourly delays in time to treatment (TTR) for severe infections increase mortality [11]. Thus, there is a pressing need for the development of novel AST methodologies that can detect all resistance mechanisms and provide rapid, precise, and consistent results to better inform antibiotic treatment strategies [8, 12]. There have been several attempts at developing such methods in the last decade with a variety of approaches [13, 14, 15, 16, 17, 18, 19, 20, 21, 22]. These rapid phenotypic methods suffer from inherent limitations that prevent them from becoming more widely adopted, including cost, reliance on databases, narrow antibiotic ranges, and high false positive rates [23, 24, 25, 26]. Agar pads are a standard format to analyse and image bacteria in optical microscopy, e.g. [27]. Of particular note for us was the work done by Choi *et al*., where they made a customized injection-moulded platform for high throughput single-cell imaging used for AST [17]. We wanted a more flexible design that is easier to fabricate, confining bacteria to a single imaging plane for improved image quality and removing the dependence on antibiotic diffusion.

To address these issues, we developed the Multipad Agarose Plate (MAP), a new platform for phenotypic AST that addresses key limitations of existing methods. The MAP is designed on a standard well-plate format and contains 96 agarose pads. Each pad serves as an independent micro-environment, where media and drugs can be added independently. We have developed reliable protocols for making and inoculating these pads. Standard brightfield illumination and timelapse microscopy enable tracking within minutes without the need for labelling or staining. We have made a fully automated analysis toolchain that performs robust segmentation to detect the microcolonies and track their growth rates. By configuring concentration gradients of antibiotics across the pads we show that the MAP can be used to perform rapid AST.

In this work, we start by demonstrating that the MAP is robust in response to variations in bacteria sample density, illumination wavelength, and agarose concentration in the pads. Then, the application of the MAP in conducting antibiotic susceptibility testing (AST) by tracking bacterial growth rates is validated. Under our experimental conditions, which included mono-cultures of *E. coli* K-12 MG1655 with nine antibiotics, we were able to obtain reliable minimum inhibitory concentrations (MICs) within a three-hour incubation period with the antibiotics. These MICs align well with those derived from control methods: broth microdilution, ETest, and data published by the European Committee on Antimicrobial Susceptibility Testing (EUCAST).

## 2 Materials and Methods

### 2.1 Preparation of MAP platforms

Each platform was assembled using pre-cut sheets of acrylic and adhesive. The base and well plates were laser-cut from 5 mm and 1.5 mm cast acrylic sheets, respectively (RS Components (UK), 082-4676 and 082-4480). The two adhesive sheets were laser-cut from 3M 468 sheets with thickness 0.13 mm (Self Adhesive Supplies (UK), 113317).

The pads are made by mixing the desired combination of growth media and any temperature-stable constituents with 1% w/v agarose powder (Sigma-Aldrich, A9539). The agarose mix is then heated to 80 °C to fully dissolve the agarose powder and cooled to 60 °C before adding temperature-sensitive constituents such as antibiotics. 31 μL of the mix is then transferred to each MAP well, where the agarose is allowed to solidify at room temperature. We used an Opentrons OT2 pipetting robot to prepare dilution series of antibiotics and make the pads on the MAP platform using an 8-channel P300 OT-2 electronic pipette [28]. At this stage, the MAP is ready for seeding the samples. The platforms can be made in batch, lidded and stored at 4 °C. We have verified the shelf-life of a ready-made platform to exceed three days. For example, repeats 1 and 2 of our datasets for ampicillin and vancomycin (as shown in fig. S6.H and I respectively) were collected on the same day as the MAP pads were made, whereas repeats 3 and 4 were collected three days later; no significant differences are observable.

Making the agarose pads is the most challenging aspect of using the MAP. If the pads are made by adding too much volume, fluid from the pads can move between the pads when confining the bacteria sample with the glass slide. Equally problematic, if the wells are filled with too little agarose media, the pads will not contact the glass slide, rendering the pads impossible to image. To balance this, we found it best to fill the wells with a volume of agarose media that was just above the well capacity and wait for five to fifteen minutes, allowing the pads to evaporate to just the right size before adding the lid and refrigerating.

The acrylic and adhesive sheets were clean but not sterile in our experiments. Similarly, the pads were made from sterilised media in a non-sterile environment. In practice, we find that the concentration of contaminants ends up being negligible compared to that of the seeded samples. For special applications, the MAP platforms can be sterilised with Ultraviolet Germicidal Irradiation (UVGI) and the pads can be made in a sterile environment.

### 2.2 Strains, growth conditions and seeding

In all experiments, *E. coli* K-12 strain MG1655 with LB broth (ThermoFisher, 10855001, with 10 g peptone, 5 g yeast extract, 5 g sodium chloride per 1 L media) as growth media was used. Pre-cultures were grown overnight and diluted 500x into fresh media and allowed to resume exponential growth for at least three generations. These seed cultures were then diluted to an optical density at 600 nm (OD600) of 0.02, and aliquots of 1.5 μL were placed on the surface of each MAP pad. The pads were then dried for 5 to 10 minutes before peeling the protective film on the upper adhesive sheet and sealing the pads with a single glass slide that covers all the pads (UQG Optics, GPD-1577, dimensions 110 × 74 × 0.17 mm).

### 2.3 Timelapse microscopy

The MAP platform was then imaged at 37 °C on a custom-built, inverted microscope for 4 hours. We imaged one field of view (FOV) per pad, using a looping script to capture all the images automatically. A LED focusing system was used to keep the sample roughly in focus automatically. To account for possible misalignment between the camera’s focal plane and the imaging plane of individual pads (which can exhibit significant tilts in relation to each other on the same MAP), a z-stack of images was captured for each FOV. This approach ensures that each microcolony within the frame is captured at its optimal focus, despite the spatial variations in the imaging planes across different pads. For these experiments, we chose the FOVs manually to optimize the quality of our data sets. However, selecting FOVs could be done in an automated fashion to give a fully automated imaging workflow.

Imaging was performed with brightfield illumination using the Nikon 40x CFI Plain Flour air objective with a numeric aperture of 0.75. The camera was a Teledyne FLIR BFS-U3-70S7M-C with a 7.1 MP Sony IMX428 monochrome image sensor. All images were captured at 3208 × 2200 pixels, resulting in an effective resolution of 0.112 μm*/*pixel. The temporal resolution of the datasets is 8 to 12 minutes, limited by the time it takes for the microscope to image all pads.

### 2.4 Antibiotics

The antibiotics we used to mix into the pads of the MAP were ampicillin (Sigma-Aldrich 10835242001), carbenicillin (Sigma-Aldrich C1389), ciprofloxacin (Sigma-Aldrich 17850), chloramphenicol (Sigma-Aldrich C0378), kanamycin (Sigma-Aldrich K4000), mecillinam (Sigma-Aldrich 33447), tetracycline (Sigma-Aldrich T3258), rifampicin (Sigma-Aldrich 557303) and vancomycin (Sigma-Aldrich V2002). Stock solutions of 50 g L^*−*1^ were prepared by dissolving the antibiotic in 10 mL of milliQ water (ampicillin, carbenicillin, ciprofloxacin, kanamycin, mecillinam, tetracycline, vancomycin), 100% methanol (rifampicin), or 95% ethanol (chloramphenicol) and stored at *−*20 ^*°*^C until use.

### 2.5 Image processing

The images are analysed using a customised processing pipeline available in *PadAnalyser*. It works as follows:

#### Z-stack projection

Each z-stack is projected into a single frame using a focus stitching algorithm. Each frame in the stack is subdivided into tiles of 200 × 200 px, and the focus score is computed as the sum of the square of the Laplacian transform. For each stack, the frame with the highest score is kept to make up the in-focus image, fig. S3.

#### Normlization

Each projected frame is normalized by clipping the brightest and darkest pixels, applying a Gaussian blur with 3x3 kernel, and normalizing the result to integers in the range 0 to 255. These frames are stored as 8-bit grayscale images.

#### Colony segmentation

Colonies are segmented using the Scikit Image Canny Edge Detector [29] with *σ* = 1, and performing a morphological close with a circular kernel of size 7 × 7 and two iterations. This produces a binary image with masks representing the colonies. For details, see fig. S2.

#### Colony filtering

The detected masks are then filtered to remove any masks that are within 20 pixels of the edge of the frame or have an area less than 2 μm. Debris and out-of-focus masks are removed by computing the Laplacian intensity profile of each mask as a function of distance from the mask edge. For actual colonies, this profile will have a positive peak within 5 pixels, and a negative peak within 10 pixels of the mask edge. Masks that do not satisfy this criterion are removed. For details, see fig. S2.

#### Frame alignment

The centroids of the segmented colonies are then used to align consecutive frames to compensate for thermal expansion and stage drift.

#### Linking colonies over time

Next, colonies from the different time steps of a FOV are linked and assigned an identity. For details, see fig. S2.

#### Extracting statistics

Properties including area, arc length, centroid and lineage are computed and exported for each colony for each timestep. Based on this data, colony growth rates are computed based on the rate of change in colony area over time using the *gaussianprocessderivatives* python package [30].

#### Debug output

Log and video files are produced for each field of view with segmentation mask outlines drawn in clear colours so the user can validate that the algorithms are working correctly.

For the colony area to be a good proxy for colony biomass, this method assumes that the bacteria spread out in a single layer. We observe that the colonies grow in a single layer until they have formed a very dense colony with hundreds of bacteria, consistent with our previous work [31]. For our *E. coli* strain in LB broth, a transistion to stacking (growing into more complex 3D structures) occurs after approximately three hours. Further details can be found in section 3.1.

### 2.6 Data fitting

AST curves were fitted to growth rate data using Hill functions

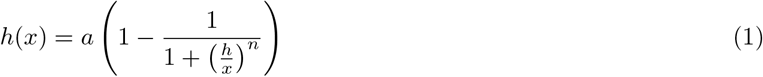

where *a, h*, and *n* are fitting parameters. Based on these fits, inhibitory concentration 90% (IC_90_) were obtained by 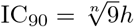 [32].

### 2.7 Broth microdilution experiments

Microplates with 384 wells were set up with the same growth media, antibiotic stocks, sample preparation protocol and bacteria strain used for the MAP experiments. Growth media and antibiotics were pipetted into each well before adding bacteria to produce an effective OD600 of about 0.05. The plate was then incubated at 37 °C with OD measurements every 20 minutes on a BMG FLUOstar Omega Microplate Reader.

We chose the Minimum Inhibitory Concentration (MIC) as the lowest antibiotic concentration capable of inhibiting OD600 by at least 80% relative to the growth control after 16 hours of incubation. This selection criterion aligns with the European Committee on Antimicrobial Susceptibility Testing (EUCAST) method for broth microdilution. While the standard EUCAST method relies on visual inspection of wells showing no growth, this quantitative approach is particularly recommended for antibiotics such as trimethoprim and trimethoprim-sulfamethoxazole [9].

### 2.8 ETest experiments

The ETest assays were performed using identical *E. coli* cultures to those used in MAP and broth microdilution experiments. Precultures were diluted to an OD600 of approximately 0.2 prior to plating on standard LB agar plates (90 mm). The resulting bacteria lawn was dried before two ETest strips were placed on each plate. After incubating for 24 hours at 37 °C, the plates were examined and the MIC was determined based on the scale on the strip, adhering to the Biomerieux reading guide [10]. The specific ETest strips used included ampicillin (Biomerieux 412253), chloramphenicol (Biomerieux 412309), ciprofloxacin (Biomerieux 412311), kanamycin (Biomerieux 412382), mecillinam (Liofilchem 920171), rifampicin (Biomerieux 412450), tetracycline (Biomerieux 412471), and vancomycin (Biomerieux 412488). Note that ETest strips for carbenicillin were not obtained for these assays.

## 3 Results

The process of using the MAP platform is summarized in fig. 1. The MAP platform, shown in Figure 2.A, consists of 96 agarose pads arranged in a 12 × 8 grid. It is simple to assemble from a glass slide and two laser-cut sheets of acrylic and adhesive, as shown in the cross-sectional view in fig. 2.B. Detailed instructions for assembling and using the platforms are in Materials and Methods. Each pad provides a separate growth environment where media and drugs can be added independently (see section MAP supports consistent growth and imaging of microbes). After loading the bacteria samples on each pad, the platform can be imaged with time-lapse optical microscopy. A fully automated segmentation pipeline analyzes the microscopy data and tracks statistics about the colony-forming units (CFUs) and how they grow into microcolonies. Figure 2.C shows cropped frames at varying points in time where *E. coli* K-12 MG1655 is growing in LB broth. The frames are annotated with their segmentation masks.

**Figure 1:**
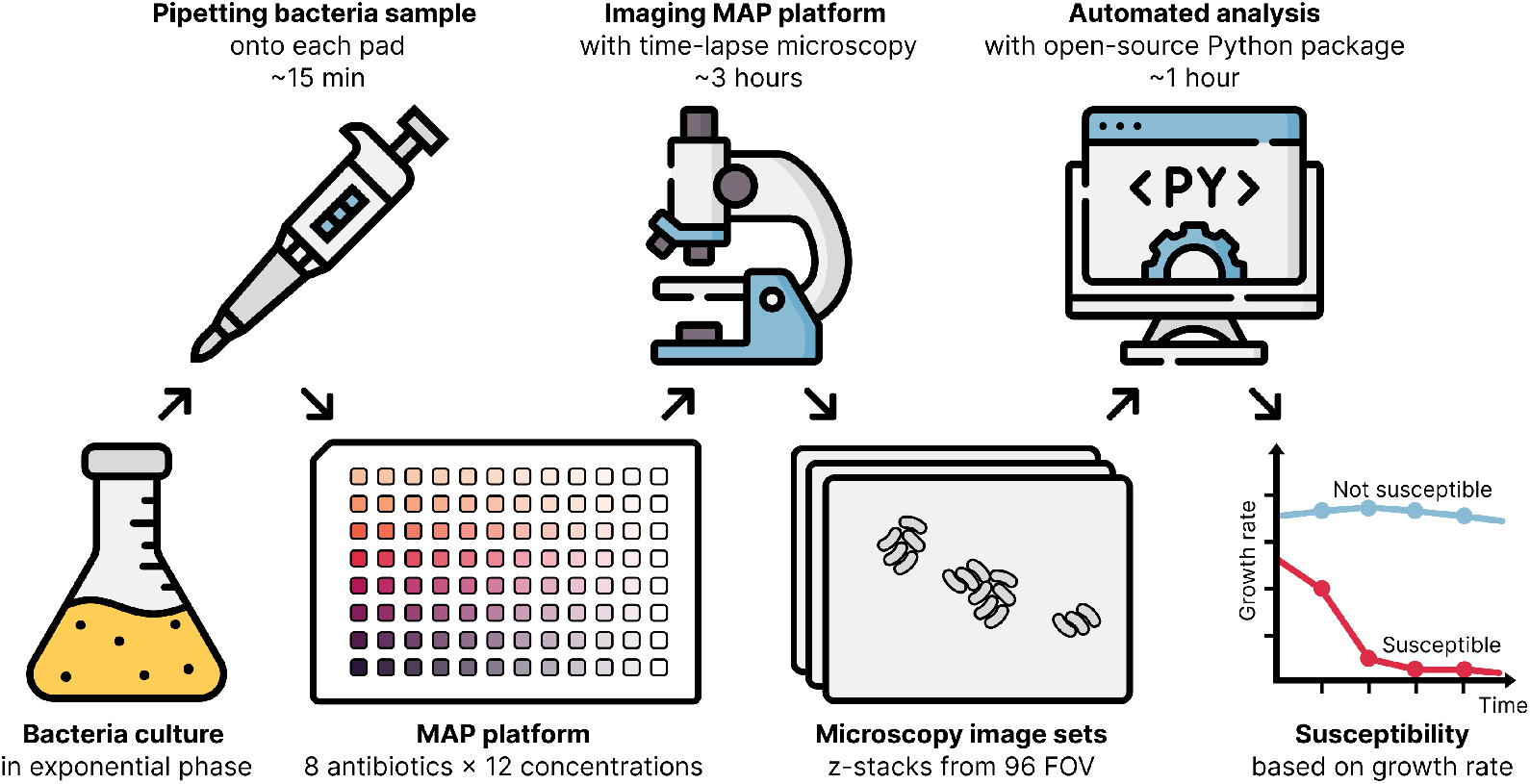
An overview of how to use the MAP for AST. A bacteria culture in the exponential phase is pipetted onto the MAP platform that has been prepared with pads containing eight antibiotic dilution series. The platform is then imaged with brightfield microscopy. The resulting images are processed with a fully automated analysis pipeline that segments the microcolonies and tracks their growth rates. The growth rates are then used to determine the susceptibility and the minimum inhibitory concentration (MIC) of the antibiotics.

**Figure 2:**
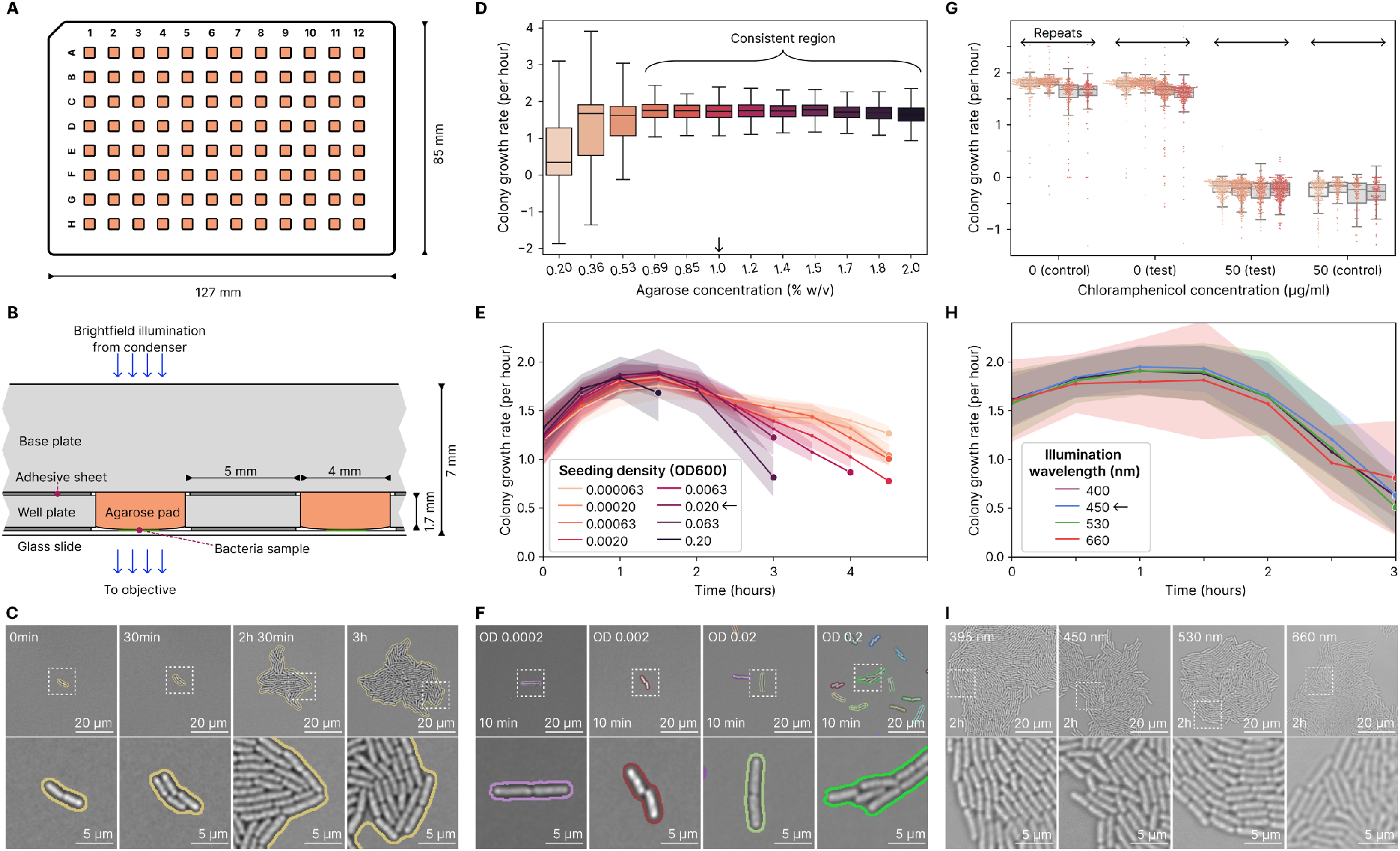
The MAP platform can be used to measure bacteria growth automatically with high resolution and reproducibility. **A**: Schematic representation of the MAP. The platform features 96 square pads of 4 mm size, arranged in a 12 × 8 grid with 9 mm pitch. The standard well-plate format facilitates compatibility with standard multichannel pipettes and stage holders. **B**: A detailed cross-sectional view of the MAP illustrates the path of brightfield illumination. The light traverses through the base plate and agarose pad before intersecting with the imaging plane. Like most biology labs, we use inverted microscopy, where the objective is located below the sample. The acrylic base and well plates are kept together using the same design of adhesive sheet that glues the glass slide to the well plate. The bacteria sample grows in the interface between the agarose pad and the glass slide. **C**: Cropped frames with time-series microcolony growth of *E. coli* on LB broth with 1% w/v agarose at 37 °C. This sequence of images illustrates the expansion of a colony-forming unit (CFU) into a microcolony. The border marks the colony segmentation masks. Time is reported as time after imaging is started, which is typically 20 to 30 minutes after the bacteria are placed on the pads. After 2.5 hours of growth, the microcolony is still growing in a single layer, but half an hour later, stacking has started to occur. **D**: Varying the agarose concentration does not significantly affect the growth rate between 0.069 to 2% w/v. For low agarose concentrations, growth rates cannot be consistently tracked, leading to very large standard deviations. The data represents twelve replicate pads for two repeat experiments. The arrow indicates how 1% was chosen as optimal. Figure S1.A shows how the MAP platform was set up. **E**: Varying the optical density (OD600) of the liquid bacteria samples placed on each pad affects the density of CFUs. Each line represents the average colony area growth rate over time after imaging is started on the microscope. The lines terminate when there are less than four colonies tracked for a given condition. The data represents seven replicate pads per seeding density. The arrow shows that an OD of 0.02 was used for further experiments. Figure S1.B shows how the MAP platform was set up. **F**: Frame sections from the first 10 minutes of imaging illustrate how seeding densities typically look on the pads. Below OD 0.002, most FOVs do not have any bacteria present. **G**: An assessment of cross-contamination between adjacent pads on the MAP shows individual pads are unaffected by their neighbours. Test pads containing 0 μg mL^*−*1^ of antibiotic were placed around pads containing 50 μg mL^*−*1^ of antibiotic, and vice versa. Each data point in the plot represents the growth rate of a single colony within the initial 3 hours after imaging was started. Each repeat contains data from about twelve pads per condition. Figure S1.C shows how the MAP platform was set up. **H**: The *E. coli* exhibit little to no variation in growth rate for different wavelengths of brightfield illumination during time-lapse imaging. 395, 450, 530, and 660 nm light was tested, corresponding to ultraviolet, blue, green, and red, respectively. The arrow shows that 450 nm was chosen as optimal. The data represents twelve replicate pads per illumination wavelength for three repeat experiments. Figure S1.D shows how the MAP platform was set up. **I**: Images are included after two hours of growth to illustrate how bacteria look when illuminated by the different wavelengths.

### 3.1 MAP supports consistent growth and imaging of microbes

*E. coli* K-12 MG1655 was used here as a model organism, because *E. coli* is an important pathogen and has the largest available literature that we can use for benchmarking. We start by conducting a set of experiments to optimize, and ensure the MAP is robust to variations in, key setup-parameters.

The concentration of agarose determines the stiffness of the pads and has been documented to affect bacterial growth [31]. Figure 2.D shows consistent growth rates between 0.69 and 2.0% agarose. Some bacteria remain motile for lower concentrations, producing large growth rate variance as the analysis tool struggles to track lineages. We chose 1% agarose as the optimal concentration for subsequent experiments, as lower agarose concentrations reduce viscosity during handling and enhance pipetting precision.

The spatial distribution of CFUs on the pads will impact the number of tracked colonies and the frequency of colony merging. The concentration of bacteria in the seeding medium, combined with the volume pipetted onto each pad, determines the density of CFUs on each pad. Holding pipetted volume constant at 1.5 μL*/*pad, we varied the optical density (OD) of the seeding medium to see how this would impact tracking and growth rates. Figure 2.F shows that the bacteria generally distribute well across the pad surfaces, and for seeding densities below OD 0.002, there is (on average) less than one CFU per field of view. Figure 2.E shows that tracking with higher seeding density stops earlier, as the colonies merge and extend beyond the camera’s field of view. Low seeding densities enable tracking for the longest but place few CFUs in each field of view, so we choose an OD600 of 0.02 as optimal for further imaging. On average, this places tens of CFUs in each field of view while allowing us to track growth for three hours. See Image processing for details.

The colony growth rates change over time for all seeding ODs, as shown in fig. 2.E, following the trend expected for bacterial population growth. For all of the densities tested, bacteria grow slower at the beginning, the growth rate reaches a maximum between 1 and 2 hours after seeding and then starts to decline. The initial lower growth rate is due to adaptation, with bacteria that, having been exposed to room temperature before seeding, gear up their gene expression for growth on pads at 37 °C. After the adaptation phase, bacteria reach maximum growth rates ranging from 1.6 to 1.8 per hour, which translates to doubling times of 23 to 26 minutes, in good agreement with the 20 to 24 minutes reported in the literature for the same strain in LB broth at 37 °C [33, 34, 35]. After 2 hours, as the resources provided by the pads start to run out, growth rates start to decline, with more severe slopes corresponding to higher initial seeding densities.

The possibility of media leakage between the pads was tested. The leakage test consists of setting up MAPs with some pads containing no antibiotic and some pads with a highly effective dose of antibiotic (50 μg mL^*−*1^ chloramphenicol). These pads were arranged alternately in the central test region and clustered in the control regions (as shown in fig. S1.D). In fig. 2.G, we see that the growth rates in the control region match the growth rates in the test region, indicating no leakage of antibiotics between neighbouring pads.

Finally we explored using a shorter wavelength for the brightfield illumination during timelapse microscopy imaging to shift the diffraction limit and enable more detail to be captured in each image. Short wavelength illumination is, however, known to be more phototoxic to the cells [36]. At the intensities of our setup the data collected in fig. 2.H does not show a correlation between illumination wavelength and growth rate, all the way down to ultraviolet light. This lack of sensitivity to shorter wavelengths could be due to the short exposure time and the relatively low light intensity. We therefore chose to use 450 nm blue light, as this produced high-quality, clear images that did not seem to be significantly improved by moving to 395 nm, fig. 2.I.

### 3.2 MAP enables AST in 3 hours based on growth rate

Having optimised key setup parameters and explored the MAP’s reliability, we apply the MAP for AST using nine test antibiotics and the *E. coli* K-12 MG1655 lab strain. The MAP was set up with eight antibiotics per platform, and 11 concentrations plus a negative control for each antibiotic. The starting concentration of the antibiotic dilution series was determined in each case to place the expected MIC around the middle of the concentration gradient.

In fig. 3.A and D, we demonstrate how colony areas develop over time for tetracycline and ampicillin, with panels B and E showing the corresponding growth rates. These two antibiotics are highlighted here because they represent growth dynamics for the two broad classes of antibiotics, bacteriostatic and bactericidal, respectively. While tetracycline immediately reduces the growth rate of bacteria in a fashion that is proportional to its concentration, the apparent growth rate reduction caused by ampicillin is significantly delayed. This is due to the bactericidal mechanism of ampicillin, which causes cell lysis as a result of growth defects, and therefore requires a certain amount of growth before becoming measurable at the population level [37]. That some antibiotics exhibit this strong time-dependent effect on growth rate makes it very important to choose the time window used for AST analysis carefully. For our setup, where we have monocultures of *E. coli* seeded from exponential phase, we found the optimal time window for growth-based AST measurements to be between two and three hours after starting incubation on the MAP, as indicated by the dashed boxes in fig. 3.B and E.

**Figure 3:**
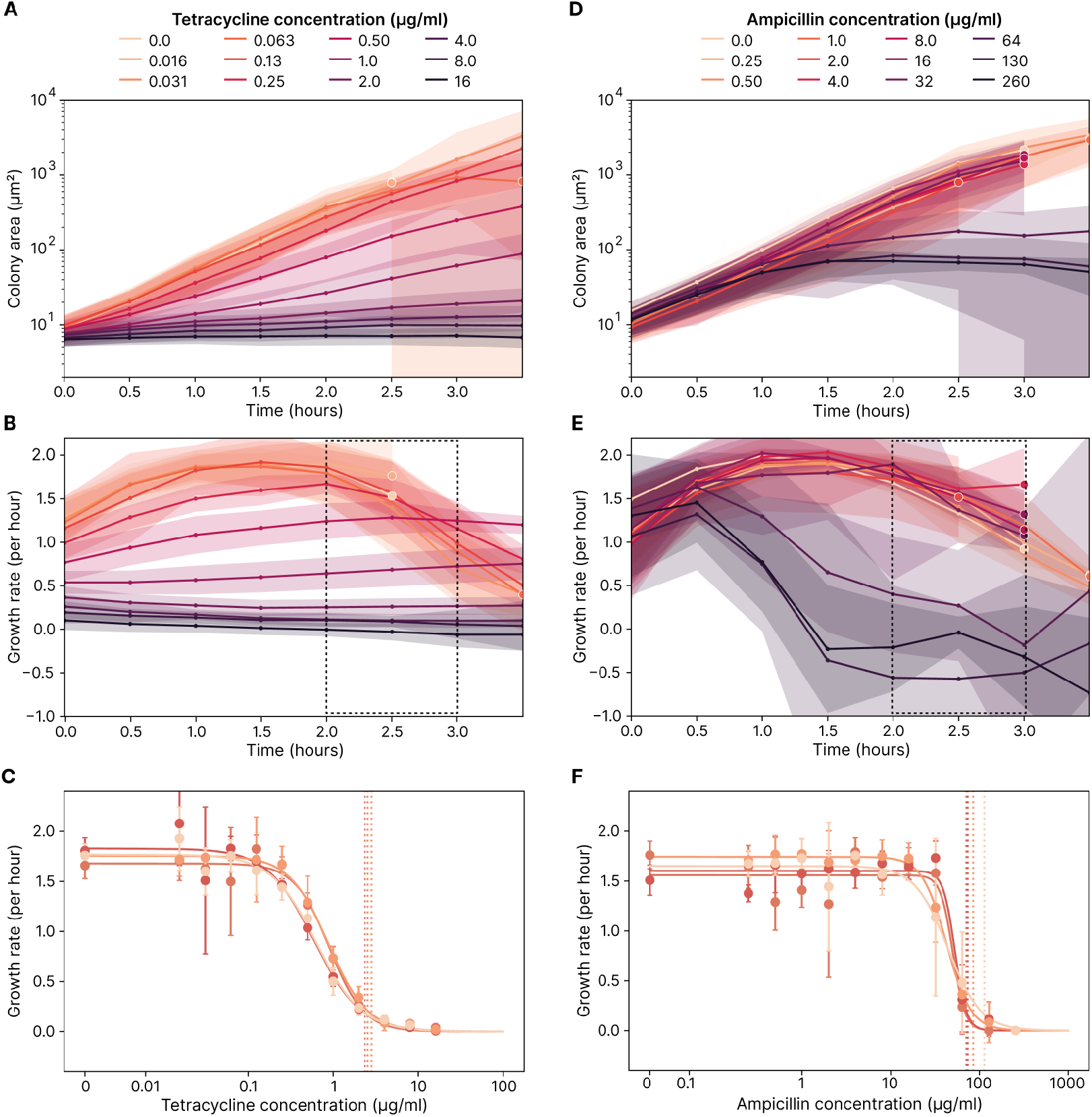
The MAP platform is used to perform AST on monocultures of *E. coli* using a test set of nine antibiotics. **A**: Colony areas develop over time for varying concentrations of tetracycline. The areas in this plot represent standard deviation, and the lines terminate in points where most colonies grow to exceed the FOV, and tracking for that pad is stopped. All experiments comprise data from four repeat experiments. **B** Colony growth rates develop over time for varying concentrations of tetracycline. Growth rates are calculated based on the time derivative of the colony area curve for each colony individually, see section 2.5 for details. The dashed growth-rate region between two and three hours indicates the time span used for evaluating AST. **C**: Growth rates are assessed for different concentrations of tetracycline. Each point in this plot is computed as the average growth rate for that given concentration in the dashed region from B, with error bars corresponding to the standard deviation. In these plots, any negative growth rate is considered spurious and set to zero. Hill curves have been independently fitted for each of four repeats, and the associated *IC*_90_ concentration is indicated with a vertical dashed line. For the four repeats, *IC*_90_ is computed to be 2.5, 2.4, 2.8 and 2.5 μg mL^*−*1^ respectively. **D**: Considering ampicillin, we see how colony areas initially develop in a similar fashion regardless of antibiotic concentration. See A for details about the plot. **E**: We see significant changes in growth rate over time for higher concentrations of ampicillin. **F**: There is a high correspondence between the four repeats with ampicillin. For the four repeats, *IC*_90_ is computed to be 71, 74, 85 and 110 μg mL^*−*1^ respectively.

The mean growth rates, between two and three hours after seeding for different concentrations of tetracycline and ampicillin, are shown in Figure 3.C and F respectively. The growth rates are fitted with a Hill curve, a sigmoidal function commonly used to quantify the effect of antibiotics on the growth rate of bacteria [32]. The *IC*_90_ point of the Hill fit (where the growth rate is inhibited by 90%) is used as MIC. Each experiment has four biological repeats with individual fits, and there is a close match between repeats.

Hill-fits from all of the antibiotics tested are summarized in fig. 4.A. As expected, there is a large range of MICs for the different antibiotics. The Hill curve fits each well, with the smallest R-squared *R*^2^ for a fit being 94% (for ampicillin) and a mean *R*^2^ of 0.98% across all fits. The complete set of area-growth curves is in fig. S5 and the complete set of Hill-fits is in fig. S6.

**Figure 4:**
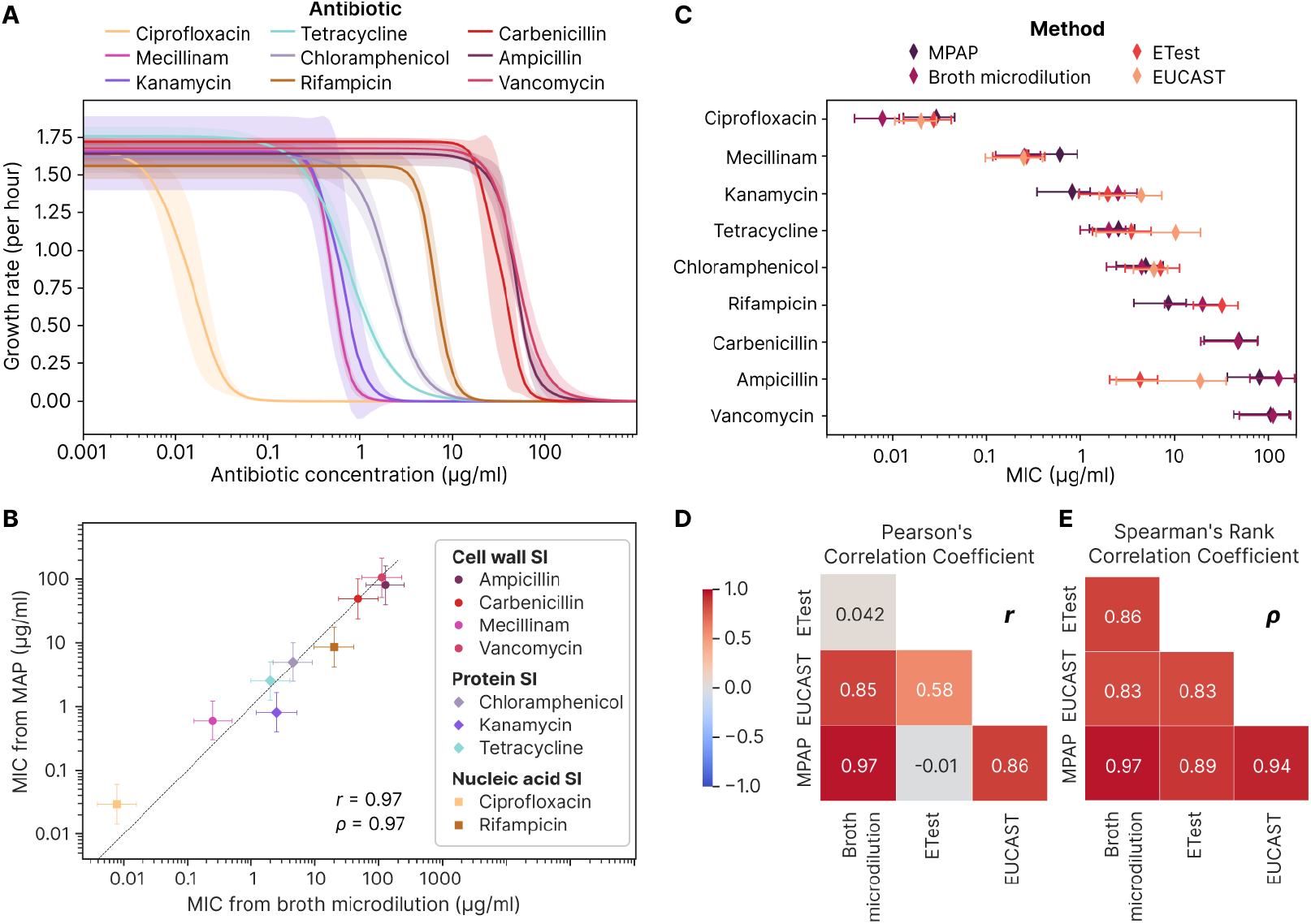
The AST results from the MAP platform correspond well with the results obtained from broth microdilution, ETest and EUCAST tabulated data. **A**: Overview of all Hill fits, with each of the nine antibiotics we evaluated. The shaded area corresponds to the standard deviation between the Hill curve fits of the four independent repeats. **B**: Comparing MIC obtained from MAP to that obtained from broth microdilution for the set of antibiotics. The dashed line is drawn for *x* = *y*. The closer to this line the data points fall, the better they correspond. **C**: Comparing MIC measured by MAP with broth microdilution and ETest comparison assays using the same strain of *E. coli* and LB growth media. Error bars are computed based on the standard deviation between repeats and the factor of two concentration steps. The EUCAST data is based on the EUCAST database of MIC distributions for wildtype *E. coli*, where the error bars represent the first and third quartile [38]. **D**: Lower triangular matrix showing Pearson’s correlation coefficient for the four methods. **E**: Lower triangular matrix showing Spearman’s rank correlation coefficient for the four methods.

The MICs produced by the MAP platform is compared with MICs from the established method of broth microdilution in Figure 4.B using the same *E. coli* cultures used for the MAP experiments. Further, we compare these results with MICs obtained from ETest strips and EUCAST tabulated data in fig. 4.C and table 1. The EUCAST data is based on the EUCAST database of MIC distributions for wildtype *E. coli*, primarily using Mueller Hinton broth as growth medium [39]. The methods produce consistent MICs for most antibiotics. Notable exceptions include carbenicillin (no data available from EUCAST and difficult to procure ETest strips) and ampicillin (which is found to be less effective based on the MAP and broth microdilution assays). In addition, we find the MG1655 strain to be susceptible to high concentrations of vancomycin using the MAP and broth microdilution methods. While vancomycin has a large structure that limits its permeation of the outer membrane of gram-negative bacteria, it has nevertheless been documented to accumulate in gram-negative bacteria at higher concentrations [40].

**Table 1:** Comparison of minimum inhibitory concentrations (MIC) for nine antibiotics: assessment by the MAP Platform versus Broth Microdilution and ETest Methods using *E. coli* K-12 MG1655. Data from EUCAST literature review for *E. coli*. Annotations: “none” denotes an unobtainable value; “≥ 254” indicates resistance to the maximum concentration in the ETest.

| Antibiotic | MIC (μg/ml) |  |  |  |
| --- | --- | --- | --- | --- |
|  | MAP | Broth microdilution | ETest | EUCAST |
| Ciprofloxacin | 0.03 ± 0.02 | 0.008 ± 0.004 | 0.03 ± 0.01 | 0.020 ± 0.009 |
| Mecillinam | 0.6 ± 0.3 | 0.3 ± 0.1 | 0.3 ± 0.2 | 0.2 ± 0.1 |
| Kanamycin | 0.8 ± 0.5 | 3 ± 2 | 2 ± 1 | 4 ± 3 |
| Tetracycline | 3 ± 1 | 2 ± 1 | 4 ± 2 | 10 ± 9 |
| Chloramphenicol | 5 ± 3 | 5 ± 3 | 7 ± 4 | 6 ± 2 |
| Rifampicin | 9 ± 5 | 20 ± 10 | 30 ± 20 | none |
| Carbenicillin | 50 ± 30 | 50 ± 30 | none | none |
| Ampicillin | 80 ± 40 | 130 ± 60 | 4 ± 2 | 20 ± 20 |
| Vancomycin | 110 ± 60 | 110 ± 60 | ≥ 254 | none |

To obtain a quantitative estimate of how well the MICs measured with these different methods correspond, we computed Pearson’s and Spearman’s rank correlation coefficient matrices in fig. 4.D and E respectively. Both Pearson’s correlation coefficient and Spearman’s rank correlation were very high for the MAP and broth microdilution MICs, indicating strong correspondence between the results. While the microdilution and MAP antibiotic stocks were prepared in-house, ETest strips are commercial and experiments for EUCAST were carried out by others with varying strains of *E. coli*, introducing a significant level of variability. Finally, the Pearson coefficient was low for the ETest - MAP and ETest - suspended dilution correlations, which could be attributed to Pearson’s sensitivity to outliers (such as the value obtained for ampicillin) since Spearman’s rank correlation remained high for these combinations.

## 4 Discussion

Rapid and accurate AST is urgently needed to improve infection treatment, combat the rise of antimicrobial resistance and preserve the shrinking pool of effective antibiotics that remain in our arsenal. While genotypic methods offer speed, they lack the comprehensive coverage of phenotypic assays, which remain the gold standard due to their direct measurement of bacterial response to antibiotics. However, the clinical adoption of rapid phenotypic AST methods is hindered by their complexity and cost. Commonly used phenotypic methods in clinical settings, such as broth microdilution and disk diffusion, are reliable but slow, typically requiring 16-48 hours to yield results. On the other hand, emerging rapid phenotypic methods, though faster, often involve intricate and expensive setups, limiting their widespread use in clinical laboratories.

The MAP (Multipad Agarose Plate) introduced in this study addresses these limitations by offering a solution that is not only rapid but also simple and cost-effective. Compared to its most successful predecessor [17], the MAP is much easier and affordable to fabricate, not requiring expensive micro-injection molds, and yields extremely high-quality microscopy images. Our demonstration that the MAP can deliver reliable AST results within a three-hour window marks a significant improvement over traditional phenotypic methods, aligning with the speeds achieved by more advanced, yet less accessible, AST systems.

The versatility of the MAP extends beyond its current application. While this study focused on *E. coli* to benchmark against established MIC values, the MAP’s design is conducive to analyzing more complex clinical samples. Its capacity to distinguish cells based on morphology makes it well-suited for poly-microbial infections and samples containing host cells. This adaptability could drastically reduce the time and complexity involved in sample processing in clinical diagnostics. Further to interspecific diversity in the samples, MAP could also be used to study intraspecific differences such as those brought about by phenotypical heterogeneity that leads to the production of persister cells that drive infection relapses. This could open the door to more nuanced, specialized treatments promising improvements in patient care and also to high-throughput investigations of fundamental microbiology.

## 5 Conclusion

MAP represents a significant advancement in the field of AST. The platform’s unique combination of speed, simplicity and cost-effectiveness, coupled with its potential for broader applications, positions it as a promising tool in both clinical and research contexts. Further development and validation of the MAP in diverse microbial contexts will enhance the platform’s utility and pave the way for its adoption in routine clinical practice, ultimately reducing the time to test results and contributing to better patient outcomes.

## Supplementary Information

Supplementary information is available for this paper with additional data and figures that provide further insight into the study.

## Acknowledgments

We thank Emma Jones, Erika Causa and Aske Petersen for insightful feedback and support. The project was funded by the EU EC ITN Phymot, Marie Sklodowska-Curie grant agreement No 955910. LM acknowledges funding from the Herchel Smith Postdoctoral Fellowship.

## Data availability statement

Our open-source Python package, PadAnalyser, can be found at github.com/Cicuta-Group/PadAnalyser and installed through the Pipy package manager. This package contains the entire analysis pipeline we use for image preprocessing, segmentation, statistics extraction and plotting. Instructions for how to make, assemble and use the MAP platform can be found at github.com/Cicuta-Group/MAP-imaging, along with an example dataset and a demo showing how to use the PadAnalyser package. The data from experiments used in this work can be found at doi.org/10.5281/zenodo.8113886.

## Declarations

AD declares to work for a company that operates in the application areas described in this work, but has no direct conflict of interest.

